# Estimating the societal disease burden of East Coast fever among rural cattle keeping households of Namwala District, Zambia

**DOI:** 10.1101/2021.01.28.428585

**Authors:** Natasha Mwila, Chisoni Mumba, Omran Salih, Karen Sichibalo, Edgar Simulundu, Katendi Changula, Simbarashe Chitanga

## Abstract

The study aimed to estimate the disease burden of East Coast fever (ECF) among rural cattle keeping households of Namwala District of Zambia using Productivity Adjusted Life Years (PALYs). We modified Disability Adjusted Life Year (DALY) equations for humans to PALYs to estimate the societal burden of animal diseases. We used a structured questionnaire to collect data on parameters that feed into PALY equations. We coded and entered data from questionnaires directly into Statistical Package of Social Sciences (IBM SPSS Version 20), and entered the estimated values of PALY parameters into Mathematical Calculus Software called Integral Calculator (https://www.integral-calculator.com/). We then used the integral calculator to calculate PALY equations, which we used to estimate the societal disease burden of ECF in cattle. PALYs calculations were done in three categories; PALYs without discounting and age weighting, PALYs with only discounting, and PALYs with discounting and age weighting.

Results revealed that the years of productivity lost by a cow, bull, and ox that suffered from ECF were estimated at 15, 10, and 15 years, respectively. In the second category, the years of productivity lost by a cow, bull, and ox were seven, six, and seven years, respectively. In the final category, the years of productivity lost by a cow, bull, and ox were five years.

ECF caused a total of 517,165 PALYs in Namwala District. The quality of life reduced in years due to disability (YLD) caused by ECF per cow, bull, and ox was 0.07, 0.07, and 0.02 percent of their life expectancy, respectively. The estimated values for the years of a lifetime lost due to mortality (YLL) caused by ECF were 35, 49, and 35 percent of the life expectancy per cow, bull, and ox. These results are important for measuring outcomes of animal health problems in terms of PALYs. The findings are helpful in future projections for the future burden of any disease and can be used as a basis in policy-making and decision-making, particularly on priorities in animal health research. We recommend that a classification of animal diseases of national economic importance should consider both the societal burden and economic impact instead of the common practice of only considering the economic impact.

## Introduction

Livestock production is an important socio-economic activity in Zambia and contributes about 1.9% to its gross domestic product (GDP) [1]. In Zambia, livestock production is broadly categorized into commercial and traditional sectors [2]. Commercial beef farmers own large herds of mostly exotic breeds of cattle and contribute 16% of the total cattle population [3]. In contrast, the traditional sector maintains the largest cattle population at 84% and consists of smallholder farmers that mostly keep local breeds of cattle integrated with crop farming [3].

The government of Zambia has targeted livestock as a critical future sustainable source of revenue and as a significant component of exports diversification agenda away from copper and towards agriculture [4]. Zambia is endowed with a vast natural resource base and a relatively small human population for potential cattle production increment. The country’s extensive grazing lands offer a clear comparative advantage over its regional neighbors and provide ample capacity for Zambia to increase its relatively low cattle density [5]. An increase in cattle production would potentially result in inadequate domestic supplies, with surplus cattle for the regional market.

However, the traditional sector with a high potential for increasing livestock production is mainly hampered by the heavy burden of diseases [2]. The traditional sector is also characterized by limited disease management, limited adoption of animal confinement, high levels of animal mortality, and low productivity [1]. Thus the production of meat and milk products from cattle is small, resulting in low output volume. Low productivity combined with high prices for both beef and dairy milk has made Zambia uncompetitive locally and regionally, with neighboring countries Namibia and Botswana that have high production levels in the African region [6].

In Zambia, the primary concern that has been renowned for its devastating impact on cattle productivity is ECF [7]. East Coast fever causes high morbidity and mortality, decreased meat and milk production, loss of draught power and manure, thereby causing significant social and economic distress to the individual farmer. East Coast fever causes significantly more deaths than the other tick-borne diseases combined [6]. Control measures have been employed by the Veterinary Department and traditional farmers, including dipping, Immunization, and treatment, which are costly [8]. Despite the efforts, ECF is still the major disease problem and constraint on Zambia’s livestock development [8]. Previous studies have estimated the financial and economic impact of the cost of control due to ECF [9][10][11]. While the financial and economic impact of ECF control has been determined, there has been no attempt to estimate how losses of cattle and their associated products (milk. meat, draught power) impact rural communities. Therefore, this study focused on estimating the societal disease burden of ECF among rural cattle keeping households by estimating both the quality of life reduced due to disability (morbidity in terms of productivity) and lifetime lost due to premature mortality using Productivity Adjusted Life Years (PALYs). The method of PALYs used for this study is unique. It estimates productivity losses of an animal for both morbidity and mortality and quantifies the analysis of a disease burden of which the results can be used to analyze a cost-effective alternative intervention.

## Materials and Methods

We obtained ethical approval from ERES Converge of Zambia with ethical clearance number (2017 – Jul – 021). Individual verbal consent was obtained from each participant through verbal explanation of the study purpose using English, Tonga, and Nyanja languages.

### Study sites and design

We used a cross-sectional study design employing a quantitative data collection technique on cattle farmers’ population in Namwala District of Zambia. Namwala District was chosen because it has the highest cattle population in Zambia, estimated at 145,704 [12] with an estimated human population of 82,810 [13].

### Sample size calculation

We used Epitools (http://epitools.ausvet.com.au/) to calculate the sample size. Given a total population of 82,810 cattle farmers [13], a confidence level of 95%, an estimated proportion of 50%, and desired precision of 5%, the necessary sample size was calculated at 385 respondents. We sampled and interviewed cattle farmers from Chitongo, Kabulamwanda, Maala, and Namwala Central veterinary camps.

### Sampling Techniques

We approached the District Veterinary Officer (DVO) to provide a list of veterinary camps accessible by road and had a large number of cattle. We, therefore, selected Chitongo, Kabulamwanda, Maala, and Namwala Central veterinary camps. Veterinary camps are the smallest administrative offices in the district and are manned by veterinary assistants who all report to the District Veterinary Officer [14]. These veterinary camps formed a sampling unit for the study. In each veterinary campsite we used a simple random sampling technique to get the required sample size from a larger population. Veterinary camps with a larger number of cattle farmers had a higher proportion.

### Data collection techniques

A structured questionnaire was developed to capture data on a wide range of variables related to the number of cattle owned per farmer, reasons for cattle keeping, health condition, cattle productivity, cattle morbidity and mortality, and cost structures on control of ECF. The field data collection was done in two seasons; in the cool and dry season and hot and dry season, to factor in other seasonal conditions. However, this did not affect the results that were obtained. The questionnaire was pretested at the Namwala Central veterinary camp to assess the clarity, practicality, feasibility, validity, and ambiguity. This was done to ensure high-quality data collection. The questionnaire was then revised after the pilot testing to improve clarity. The interviews were carried out in English, and for those farmers who were not able to communicate in English, we translated the questions to their respective dialects, including Tonga, Nyanja, and Bemba. A structured questionnaire was administered in a face-to-face interview. Farmers were interviewed at abattoirs, district veterinary office, local markets, and households. This was so because farmers left their households in the early hours of the day and headed to these places to sell cattle for a return, thus improving their livelihoods. This was also done because households are far from each other, making it practically impossible to visit the farmers at their homes.

### Data management and Statistical analysis

We coded and entered data from questionnaires directly into the Statistical Package of Social Sciences (IBM SPSS Version 20). We performed descriptive statistics of scale variables and frequencies for string variables in the Statistical Package of Social Sciences. We then used Mathematical Calculus Software called Integral Calculator (https://www.integral-calculator.com) by inserting the values of each parameter in the formulas for PALYs functions to estimate the societal disease burden of ECF.

### Model Assumptions

#### Productivity Adjusted Life Years

PALYs are a modification of Disability Adjusted Life Years (DALYs). PALYs for a disease or health condition are calculated as the sum of the years of life lost due to premature mortality (YLL) in the cattle population and the equivalent healthy years lost due to disability (YLD) for incident cases of the health condition [15]. PALYs calculations were done in three categories; without discounting and age weighting, with discounting but no age weighting, and with both discounting and age weighting. This was to factor in problems of validity and justice. We have derived the basic formulae for PALYs as shown in equation 1:

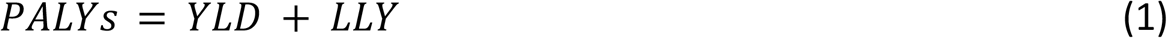

Where

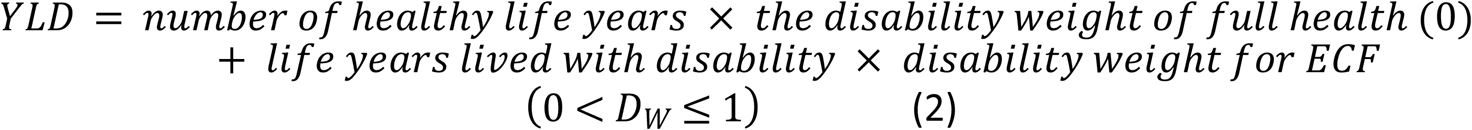

Since full health is weighted zero (0), the YLD equation reads as:

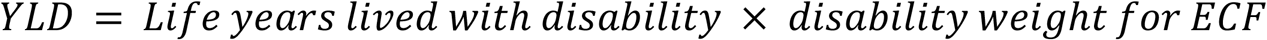

and

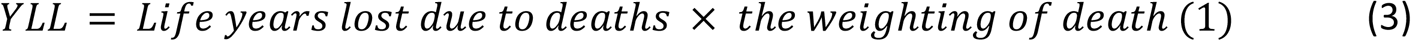

The YLD and YLL can be calculated using three methods. These include without age weighting and discounting, discounting only, and both discounting and age weighting.

#### Disability Weight

Disability is defined as some form of inability to perform everyday tasks in a usual way for cattle. Disability weight is a weight function that reflects the severity of a cattle disease between 0 (perfect health) and 1 (equivalent to death). Each disability condition is assigned a number between 0 and 1, depending on the severity of the disease [15]. Years Lost due to Disability (YLD) are calculated by multiplying the incident cases by duration and disability weight for the condition [15]. We used the disability weights developed by Salih [15], as shown in Table 1.

**Table 1:** Definition of disability weight (*D_w_*) for cattle.

| Levels | Description | $D_w$ |
| --- | --- | --- |
| 1 | 1. Beef production [(500 - 600kg for oxen), (300 - 516kg for bulls), (320 - 440 kg for cows)].<br>2. Milk production [5 - 6 litres per day].<br>3. Draught power [3 - 5hrs for cows, 5 - 6hrs for oxen].<br>4. Social status [acceptable].<br>5. Dowry payment [acceptable].<br>6. Cultural ceremonies [acceptable]. | 0 |
| 2 | 1. Beef production [(400 - 499kg for oxen), (260 - 299kg for bulls), (280 - 319kg for cows)].<br>2. Milk production [3:5 - 4:9 liters per day].<br>3. Draught power [2 - 3hrs for cows, 3 - 4hrs for oxen].<br>4. Social status [not very acceptable for the reason of loss of condition].<br>5. Dowry payment [not very acceptable for the reason of loss of condition].<br>6. Cultural ceremonies [not very acceptable for the reason of loss of condition]. | 0:01 - 0:33 |
| 3 | 1. Beef production [(300 - 399kg for oxen), (220- 259kg for bulls), (200 - 239kg for cows)].<br>2. Milk production [2- 3:4 liters per day].<br>3. Draught power [1 - 2hrs for cows, 2 - 3hrs for oxen].<br>4. Social status [not acceptable for the reason of being diseased].<br>5. Dowry payment [not acceptable for the reason of being diseased].<br>6. Cultural ceremonies [not very acceptable for the reason of being diseased]. | 0:34 - 0:66 |
| 4 | 1. Beef production [(300 - 399kg for oxen), (220 - 259kg for bulls), (200- 239kg for cows)].<br>2. Milk production [2 - 3:4 liters per day].<br>3. Draught power [1 - 2hrs for cows, 2- 3hrs for oxen].<br>4. Social status [not acceptable for the reason of being diseased].<br>5. Dowry payment [not acceptable for the reason of being diseased].<br>6. Cultural ceremonies [not very acceptable for the reason of being diseased]. | 0:67 -0:99 |

#### Discounting

Discounting means the value of a healthy life year today is set higher than the value of future healthy life years [15]. It is an economic concept that individuals prefer benefits now more than in the future. Discounting future health affects both measurements of disease burden and estimates of the cost-effectiveness of an intervention [15]. A total discounting function at any age *x* is given as indicated in equation 4.

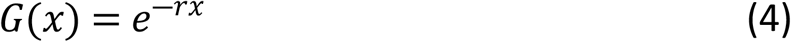

Where *r* is the discount rate

#### Age Weighting

Age weighting in DALYs means that the life years of children and older people are counted less than other ages [15]. In cattle, Age weighting determines the age at which cattle start and stop being useful in terms of milk, meat, draught power, manure, social status, dowry, and cultural ceremonies [15]. Age weighting means that cattle’s life years are counted differently because cattle are more productive at a particular age than others [15]. Therefore, in this study, we valued life experiences during productive ages based on economic and social value. The preference for productive ages is expressed mathematically, as indicated in equation 5.

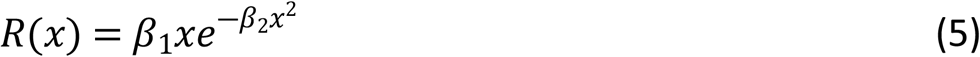

Where *x* is the cattle’s age, while *β*_1_ and *β*_2_ are parameters of the age-weighting function [15].

Figures 1 and 2 show the median age at which cattle are most and least productive for different activities. As a particular member of a cattle population grows towards the productive age, its life becomes more valuable (the age weight increase) until it reaches its maximum at the expected age of maximum productivity. Then as it gets older, its life gradually loses value [15].

**Figure 1:**
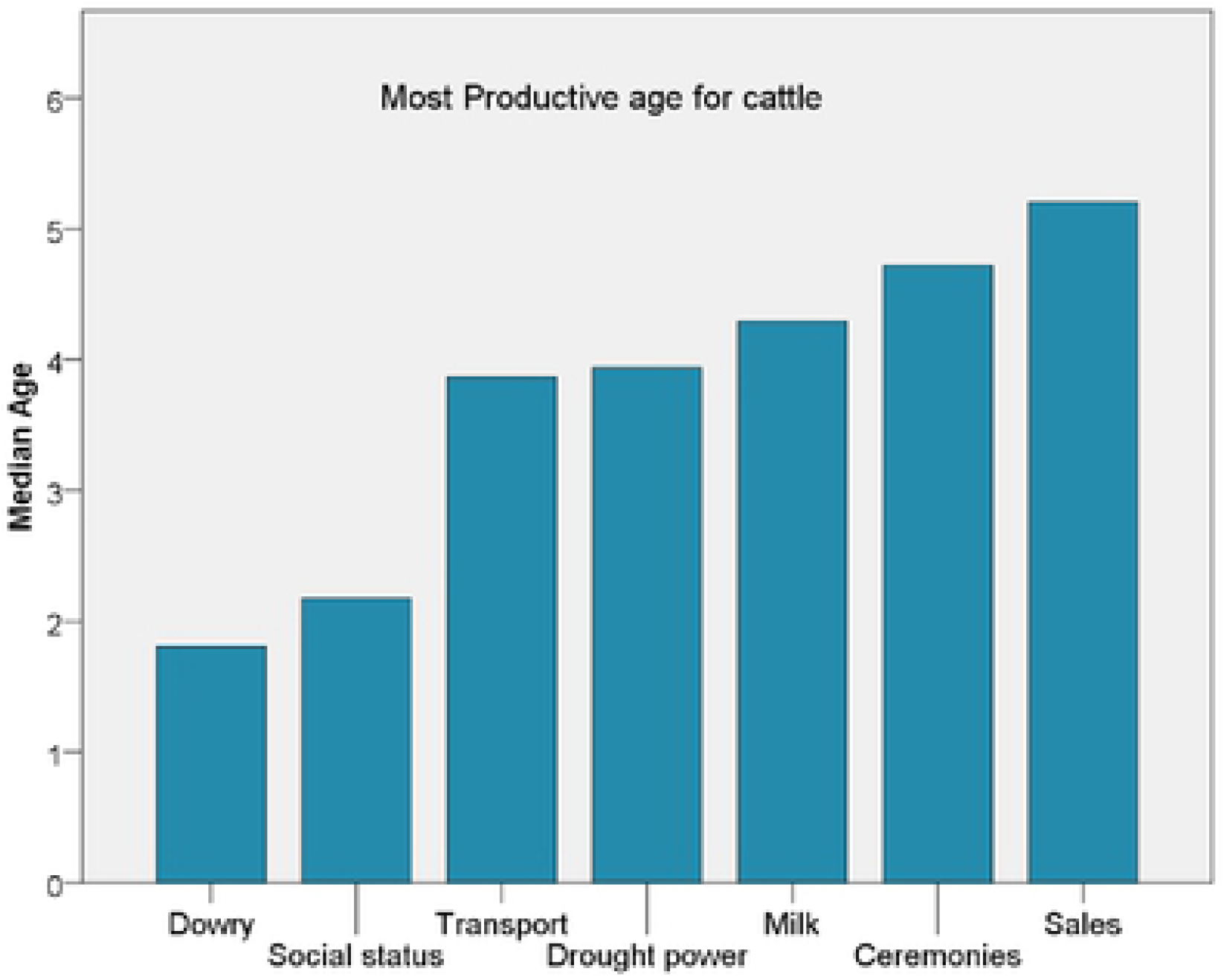
Most productive age for cattle for different activities based on questionnaires.

**Figure 2:**
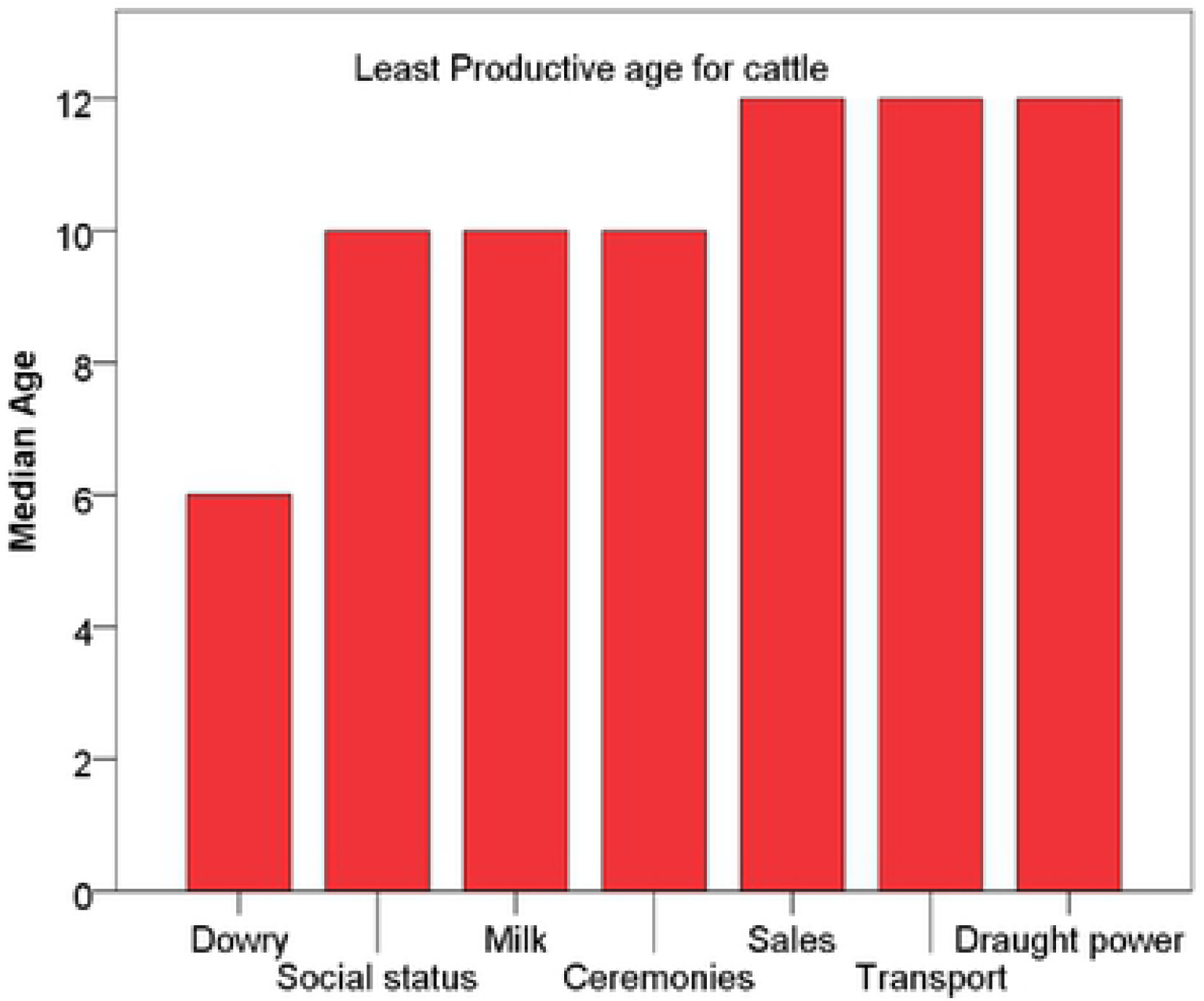
Least productive age for cattle for different activities.

#### Basic Formulas for YLL and YLD under PALYs

The basic formula for YLD (without age weighting and discounting) is the product of the number of disability cases, the average duration of the disease, and the disability weight [15]. YLD for cattle is expressed as shown in equation 6:

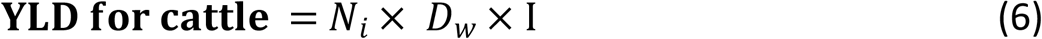

*Where N_i_ is the number of incident cases of ECF, D_w_ is the disability weight of ECF, and* I *is the average duration of the disability (ECF)*.

We used 0.5, 0.33, and 0.17 for cows, bulls, and oxen, respectively, based on each type of cattle’s different productivity roles. The basic formula for YLL (without age weighting and discounting) is defined as the product of the number of deaths and the standard life expectancy at the age of mortality. YLL is obtained, as shown in equation 7:

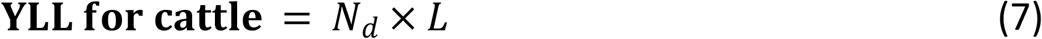

*Where N_d_ is the number of deaths, and L is the standard life expectancy at the age of death. Note that both formulae do not change whether we refer to humans or animals* [15].

#### YLD and YLL with Discounting

The second method for calculating YLD and YLL considers the discounting function. We obtained the formula for YLD by multiplying the basic YLD formula with the discounting function, as shown in equation 8.

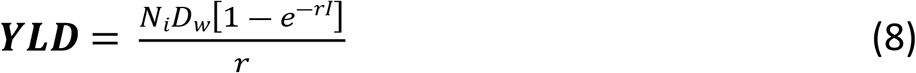

*Where N_i_ is the number of incident cases of ECF, D_w_ is the disability weight of ECF, r is the discounting rate, and* I *is the duration of the disability*.

Similarly, to find the formula for years of life lost due to premature mortality YLL, we modified equation 7 by replacing the average duration *I* by the standard life expectancy at the age of death *L*, as shown in equation 9.

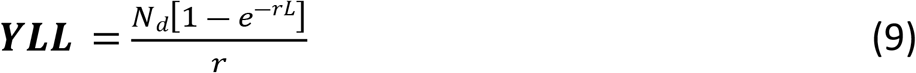

*Where N_d_ is the number of deaths, r is the discount rate, and L is the life expectancy* [15].

#### YLD and YLL with both Discounting and Age-Weighting

To calculate the YLD that accounts for the duration of the life lost due to disability (ECF), duration from the age of onset, we integrated the disability weight times the age weight and discount function over the expected period of the disability. The YLD value of any disability weight (*D_w_*) with discounting function, age weighting function, and number of disease cases (*N_i_*), as given in equation 10;

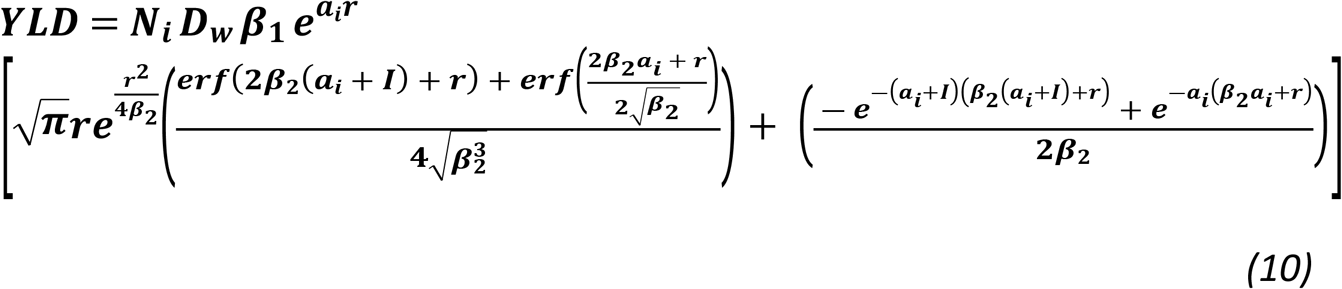

*Where N_i_ is the number of incident cases of ECF, D_w_ is the disability weight, I is the duration of disability (ECF), r is the discount rate, α_i_ is the age of onset, and erf is error function, Typical values of β*_1_ *and β*_2_ *are 0.2332 and 0.01 respectively as described by Salih* [15].

Similarly, by replacing the duration of disease *I* with the standard life expectancy *L* the age of onset with the age of death *α_d_*, we obtain the YLL formula as given in equation 11;

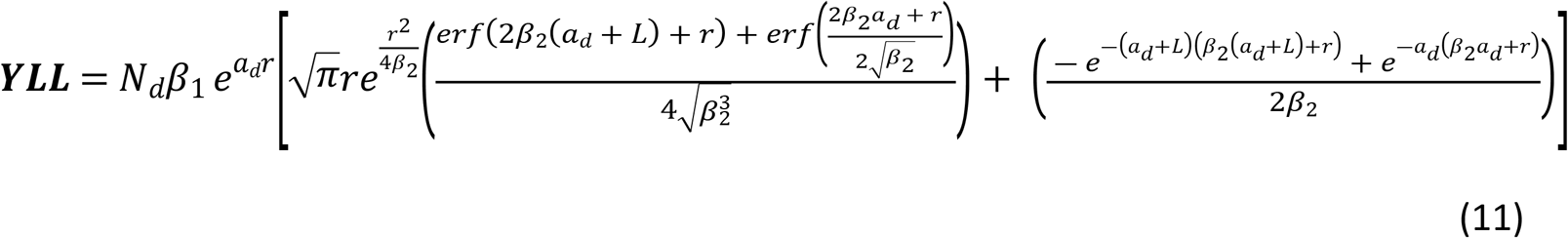

*Where N_d_ is the number of deaths, α_d_ is the age of death, and L is the standard life expectancy at the age of death*.

## RESULTS

### Estimating Societal Disease burden of ECF without discounting and age weighting

The calculations of YLD, YLL, and PALYs without discounting and age weighting for different types of cattle are shown in table 2. The YLD is 0.0096 years (approximately 1.8816 years for 196 cows), 0.0063 years (approximately 1.2617 years for 199 bulls), and 0.0033 years (approximately 0.6455 years for 198 oxen) per cow, bull, and ox, respectively. The YLL due to premature death is 14.98 years per cow (approximately 2936 years for 196 cows), 9.98 years per bull (approximately 1986 years for 199 bulls), and 14.98 years per ox (approximately 2966 years for 198 oxen). The number of PALYs lost is 14.9896 years (approximately 2937.8816 years for 196 cows), 9.9863 years (approximately 1987.2617 years for 199 bulls), and 14.9833 years (approximately 2966.6455 years for 198 oxen).

**Table 2:** Calculation for PALYs without discounting and age weighting.

| Animal | $N_i$ | $a_i$ (yrs) | $I$ (days) | $D_w$ | YLD (yrs) | $N_d$(yrs) | $L$ (yrs) | YLL (yrs.) | PALYs (yrs) |
| --- | --- | --- | --- | --- | --- | --- | --- | --- | --- |
| Cow | 196 | 4 | 7 | 0.5 | 1.8816 | 196 | 15 | 2936 | 2937.8816 |
| Bull | 199 | 4 | 7 | 0.33 | 1.2617 | 199 | 10 | 1986 | 1987.2617 |
| Oxen | 198 | 4 | 7 | 0.17 | 0.6455 | 198 | 15 | 2966 | 2966.6455 |

### Estimating Societal Disease burden of ECF with discounting

The results revealed that cattle were most affected by ECF at the age of four years, and the duration of disease was 7 days, after which an animal either responded to treatment or died of ECF. The YLD, YLL, and PALYs with discounting for cows, bulls, and oxen are shown in table 3. The YLD is 0.0096 years (approximately 1.8816 years for 196 cows), 0.0063 years (approximately 1.2597 years for 199 bulls), and 0.0033 years (approximately 1.2597 years for 198 oxen) per cow bull and ox, respectively. The YLL are 6.5979 years (approximately 1293 years for 196 cows), 5.5959 years (approximately 1114 years for 199 bulls), and 6.5979 years (approximately 1306 years for 198 oxen) per cow, bull, and ox, respectively. The PALYs lost are 6.6075 years (approximately 1.294.8796 years for 196 cows), 5.6022 years (approximately 1115.7597 for 199 bulls), and 6.6012 years (approximately 1307.2597 years for 198 oxen) per cow, bull, and oxen, respectively.

**Table 3:**
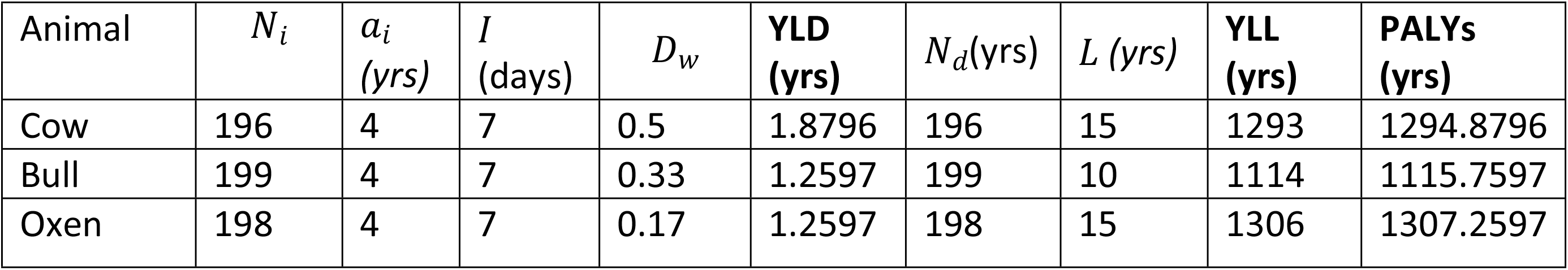
Calculation for PALYs with Discounting.

### Estimating Societal Disease burden of ECF with both discounting and age weighting

The calculation for YLD, YLL, and PALYs with both discounting and age weighting are shown in table 4. The YLD is 0.0105 years (approximately 1.9703 years for 196 cows), 0.0070 years per bull (approximately 1.3831 years for 199 bulls), and 0.0036 years per ox (approximately 0.7049 years for 198 oxen). The YLL are 5.1572 years (approximately 1011 years for 196 cows), 4.8741 years per bull (approximately 970 years for 199 bulls), and 5.1572 years per ox (approximately 1021 years for 198 oxen). The PALYs lost is 5.1677 (approximately 1012.9703 years for 196 cows), 4.8811 years (approximately 971.3831 years for 199 bulls), and 5.1608 years (approximately 1021.7049 years for 198 oxen) per cow, bull, and oxen, respectively.

**Table 4:** Calculation for PALYS with both discounting and age weighting.

| Animal | $N_i$ | $a_i$<br>(yrs) | $I$<br>(Days) | $D_w$ | YLD<br>(yrs) | $N_d$<br>(yrs) | $L$ (yrs) | YLL<br>(yrs) | PALYs<br>(yrs) |
| --- | --- | --- | --- | --- | --- | --- | --- | --- | --- |
| Cow | 196 | 4 | 7 | 0.5 | 1.9703 | 4.0192 | 15 | 1011 | 1012.9703 |
| Bull | 199 | 4 | 7 | 0.33 | 1.3831 | 4.0192 | 10 | 970 | 971.3831 |
| Oxen | 198 | 4 | 7 | 0.17 | 0.7049 | 4.0192 | 15 | 1021 | 1021.7049 |

#### PALYs Calculation on Namwala District cattle population *with both discounting and age weighting*

The total societal ECF disease burden (PALYs) for Namwala District is shown in table 5. Namwala district has a total cattle population of 145,704 [12] that comprises every cattle type. We used the herd structure developed by Lubungu [16], which states that an average of 36%, 5%, and 28% is the herd composition estimates for cows, bulls, and oxen. This translated to 52,453 cows, 7,285 bulls, and 40,797 oxen, giving us a total of 100,535 cattle population. The remaining value of 45,169 of Namwala district’s cattle population consists of calves, heifers, and steers. Using the calculation category that comprises both discounting and age weighting, ECF causes a total of 517,165.40 PALYs in Namwala District.

**Table 5:** PALYs Calculation on Namwala District Population with both discounting and age weighting.

| Cattle | Population | YLD (yrs) | YLL (yrs) | PALYs (yrs) |
| --- | --- | --- | --- | --- |
| Cows | 52,453 | 550.8 | 270,510.6 | 271,061.4 |
| Bulls | 7,285 | 51 | 35,507.9 | 35,558.8 |
| Oxen | 40,797 | 146.9 | 210,398.3 | 210,545.2 |
| **TOTAL** | **100,535** | **748.7** | **516,165.40** | **517,165.40** |

#### Calculation of PALYS with control of ECF

So far, we have calculated the PALYs lost without considering ECF control. We would like to calculate the number of PALYs that would have been averted when ECFs control is considered leading to a reduction in ECFs, i.e., if the cows were medicated. In what follows, we assume that the cows received ECF control for their disease at the age of onset (*α_i_*) or earlier, and as a result, did not die at the age of death (*α_d_*) but lived for their expected life span at the age *α_i_* (in the treated state). With these assumptions, the disability weight for the treated disease is 0.2 (in the case of a cow). Now we only need to calculate YLD with ECF control. This is achieved by changing the disability weight for the treated form of the disease from 0.5, 0.33, or 0.17 (without ECF control) to 0.2, 0.1, 0.01 (with ECF control) for a cow, Ox, and bull, respectively. In this case, the cows (and the other animals in general) would have lived for their expected life at the age of onset. In the next section, we will start with estimating PALYs using basic formulas without Discounting & Age-weighting, but with ECF control. Table 6 shows the calculation for YLD, YLL, and PALYs without both discounting and age-weighting, with age weighting only, and with both age weighting and discounting with ECF control.

**Table 6:** PALYs calculations with ECF control.

| **PALYs Calculation using basic formula (with ECF control)** |  |  |  |  |  |  |  |
| --- | --- | --- | --- | --- | --- | --- | --- |
| **Cattle** | $N_d$ | $a_i$ | $I$<br>(days) | $D_w$ | **YLD** | **YLL** | **PALYs** |
| Cow | 196 | 4 | 7 | 0.2 | Total (yrs) = 0.7518 yrs($\approx 0.003836$ years per Cow)<br>Total (days) = 275 days ($\approx 1.4$ days per Cow) | 0 | 0.7518 yrs<br>275 days |
| Bull | 199 | 4 | 7 | 0.1 | Total (yrs) = 0.3816 yrs ($\approx 0.001918$ years per Bull)<br>Total (days) = 139 days ($\approx 0.7000$ days per Bull [Less than one day]) | 0 | 0.3816 yrs<br>139 days |
| Oxen | 198 | 4 | 7 | 0.01 | Total (yrs) = 0.0380 yrs ($\approx 0.000192$ years per Bull)<br>Total (days) = 13.8600 days ($\approx 0.0700$ days per Bull [less than one day]) | 0 | 0.0380 yrs<br>13.8600 days |
| **PALYs Calculation With Age Discounting (with ECF control)** |  |  |  |  |  |  |  |
| Cow | 196 | 0.13 | 7 | 0.2 | Total (yrs) = 0.7508 yrs ($\approx 0.003831$ yrs per Cow)<br>Total (days) = 180 days ($\approx 0.9192$ days per Cow) | 0 | 0.7508 yrs<br>180 days |
| Bull | 199 | 0.13 | 7 | 0.1 | Total (yrs) = 0.3812 yrs ($\approx 0.001915$ yrs per Bull)<br>Total (days) = 91 days ($\approx 0.4596$ day per bull [less than one day]) | 0 | 0.3812 yrs<br>91 days |
| Oxen | 198 | 0.13 | 7 | 0.01 | Total (yrs) = 0.0379 yrs ($\approx 0.000192$ yrs per Ox)<br>Total (days) = 9 days ($\approx 0.0460$ day per ox [less than one day]) | 0 | 0.0379 yrs<br>9 days |
| **PALYs Calculation With both Ages Weighing and Discounting (with ECF control)** |  |  |  |  |  |  |  |
| Cow | 196 | 4 | 7 | 0.5 | Total (yrs) = 0.3552 yrs ($\approx 0.001813$ yrs per Cow)<br>Total (days) = 130 days ($\approx 0.6616$ day per cow [less than one day]) | 0 | 0.3552 yrs<br>130 days |
| Bull | 199 | 4 | 7 | 0.33 | Total (yrs) = 0.1804 yrs ($\approx 0.000906$ yrs per bull)<br>Total (days) = 66 days ($\approx 0.3308$ day per bull [less than one day]) | 0 | 0.1804 yrs<br>66 days |
| Oxen | 198 | 4 | 7 | 0.17 | Total (yrs) = 0.0179 yrs ($\approx 0.000091$ yrs per Oxen)<br>Total (days) = 6.5 days ($\approx 0.0330$ day per oxen [less than one days]) | 0 | 0.0179 yrs<br>6.5 days |

For YLD, YLL, and PALYs without both discounting and age weighting but with ECF control, the number of years of life lived with disability (YLD) is 0.7518 years per cow. The years of life lost due to premature mortality (YLL) are zero years per cow. Years of life lost due to death is equal to zero on the assumption that ECF control will avert death hence no premature mortality. Therefore, the number of PALYs lost per cow is 0.7518 (approximately 147.3528 years for 196 cows). For the bulls, the number of YLD is 0.3816 years per bull. The number of YLL is zero years per bull, as explained earlier. The number of PALYs lost per bull is 0.3816 years (approximately 75.9384 years for 199 bulls). For the oxen, the number of YLD is 0.0380 years per bull. The number of YLL is zero years per ox. The number of PALYs lost per ox is 0.0380 years (approximately 7.524 years for 198 oxen).

For the calculation for YLD, YLL, and PALYs with age weighting (with ECF control), the number of YLD is 0.7508 years per cow. The YLL are zero years per cow, as we earlier explained. Therefore, the number of PALYs lost per cow is 0.7508 (approximately 147.1568 years for 196 cows). For the bulls, the number of YLD is 0.3812 years per bull. The number of YLL is zero years per bull. The number of PALYs lost per bull is 0.3812 years (approximately 75.8588 years for 199 bulls). For the oxen, the number of YLD is 0.0380 years per bull. The number of YLL is zero years per ox. The number of PALYs lost per ox is 0.0379years (approximately 7.5042 years for 198 oxen). For the calculation for YLD, YLL, and PALYs with both Ages Weighing and Discounting (with ECF control), the number of YLD is 0.3552 years per cow. The YLL are zero years per cow, as we earlier explained. Therefore, the number of PALYs lost per cow is 0.3552 (approximately 69.6192 years for 196 cows). For the bulls, the number of YLD is 0.1804 years per bull. The number of YLL is zero years per bull. The number of PALYs lost per bull is 0.1804 years (approximately 35.8996 years for 199 bulls). For the oxen, the number of YLD is 0.0330 years per bull. The number of YLL is zero years per ox. The number of PALYs lost per ox is 0.0330 years (approximately 6.534 years for 198 oxen).

#### Calculation of PALYs averted

Having calculated PALYs without and with ECF control, now we calculate PALYs averted per cow, bull, and ox by subtracting the value of PALYs with control to those without. Table 7 shows the PALYs averted due to ECF control using the basic formula, with discounting only and with both age weighting and discounting.

**Table 7.** PALYs averted per cow, bull, and ox.

| **Calculation of PALYs averted per cow, bull, ox (Basic formula)** |  |  |  |
| --- | --- | --- | --- |
| **Cattle** | **PALYs (without ECF control)** | **PALYs (with ECF control)** | **PALYs averted** |
| Cow | 14.9892 yrs | 0.003836 yrs ( ≈ 34 hrs) | 14.9854 yrs |
| Bull | 9.9862 yrs | 0.001918 yrs ( ≈ 17 hrs) | 9.9843 yrs |
| Oxen | 14.9831 yrs | 0.000192 yrs ( ≈ 2 hrs) | 14.9829 yrs |
| **Calculation of PALYs averted per cow, bull, ox (with discounting)** |  |  |  |
| Cow | 6.6065 yrs | 0.003831 yrs ( ≈ 33 hrs) | 6.6027 yrs |
| Bull | 5.6068 yrs | 0.001915 yrs ( ≈ 16 hrs) | 5.6049 yrs |
| Oxen | 6.6023 yrs | 0.000192 yrs ( ≈ 1 hrs) | 6.6021 yrs |
| **Calculation of PALYs averted per cow, bull, ox (Ages weighting and discounting)** |  |  |  |
| Cow | 5.1682 yrs | 0.001813 yrs ( ≈ 16 hrs) | 5.1664 yrs |
| Bull | 4.8813 yrs | 0.000906 yrs ( ≈ 8 hrs) | 4.8804 yrs |
| Oxen | 5.1601 yrs | 0.000091 yrs ( ≈ 1 hrs) | 5.1600 yrs |

## DISCUSSION

This study aimed to estimate the disease burden of ECF among cattle keeping households in Namwala District of Zambia using PALYs. The results revealed that ECF causes a total loss of 517,165.40 quality years lived of cattle due to morbidity and the loss of productivity due to premature mortality regardless of the administration of treatment. The products that are expected to be lost during these years are milk, meat, manure, and draught power. A significant value of years of a healthy life is lost in livestock when ECF control measures are not put in place. On the other hand, when control measures exist, the PALYs “significantly” reduce with possibilities of increased livestock productivity. This is true regardless of the type of livestock or the method used for calculating PALYs (categories).

In the calculation for PALYs without discounting and age-weighting, the years of life lost due to disability (YLD) for a cow was 0.0096 years (approximately four days). This means that for a cow that develops ECF at the age of 4 regardless of treatment, the quality of life lived reduces by approximately 4 days. The years of life lost due to premature mortality (YLL) for a cow that dies at the age of 4 years was 14.98 years, which means that the cow losses 15 years of productivity due to ECF. The years of life lost due to disability (YLD) for a bull was 0.0063 years (approximately two days), meaning that a bull that develops ECF at the age of 4 regardless of treatment; the quality of life lived reduces by two days. The years of life lost due to premature mortality (YLL) for a bull were 9.98 years. Therefore, per bull, approximately ten years of productivity are lost due to ECF. The years of life lost due to disability (YLD) for an ox was 0.0033 years (approximately one day), meaning that an ox that develops ECF at the age of 4 regardless of treatment; the quality of life lived reduces by a day. The years of life lost due to premature mortality (YLL) for oxen were 14.98 years. Therefore, per oxen, approximately 15 years of productivity are lost due to ECF.

To calculate PALYs with only discounting, the years of life lost due to disability (YLD) for a cow was 0.0096 years (approximately four days). This means that for a cow that develops ECF at the age of 4 years regardless of treatment, the quality of life lived reduces by approximately four days. The years of life lost due to premature mortality (YLL) for a cow that dies at the age of 4 years was 6.5979 years, which means that the cow losses seven years of productivity due to ECF. The years of life lost due to disability (YLD) for a bull was 0.0063 years (approximately two days), meaning that a bull that develops ECF at the age of 4 regardless of treatment; the quality of life lived reduces by two days. The years of life lost due to premature mortality (YLL) for a bull were 5.5959 years. Therefore, per bull, approximately six years of productivity are lost due to ECF. The years of life lost due to disability (YLD) for an ox was 0.0033 years (about one day), meaning that an ox that develops ECF at the age of 4 years regardless of treatment; the quality of life lived reduces by a day. The years of life lost due to premature mortality (YLL) for oxen were 6.5779 years. Therefore, per ox, approximately seven years of productivity are lost due to ECF.

Calculation of PALYs with both discounting and age-weighting the years of life lost due to disability (YLD) for a cow was 0.0105 years (approximately four days). This means that for a cow that develops ECF at the age of 4, the quality of life lived reduces by approximately four days regardless of treatment. The years of life lost due to premature mortality (YLL) for a cow that dies at the age of 4 years was 5.1572 years, which means that the cow losses five years of productivity due to ECF. The years of life lost due to disability (YLD) for a bull was 0.0070 years (approximately three days), meaning that a bull that develops ECF at the age of 4 years regardless of treatment; the quality of life lived reduces by three days. The years of life lost due to premature mortality (YLL) for a bull were 4.8741 years. Therefore, per bull, approximately five years of productivity are lost due to ECF. The years of life lost due to disability (YLD) for oxen was 0.0036 years (approximately one day), meaning that an ox that develops ECF at the age of 4 years regardless of treatment; the quality of life lived reduces by a day. The years of life lost due to premature mortality (YLL) for oxen were 5.1572 years. Therefore, per ox, approximately five years of productivity are lost due to ECF. We incorporated both age-weighting and discounting to acquire more effective and accurate PALYs results that assessed both social values. The results without both discounting and age-weighting yielded a total of 7,898 PALYS, with discounting but no age-weighting resulted in a total of 3,718 PALYs and with both discounting and age-weighting yielded a total of 3006 PALYs. Discounting is included to prevent giving excessive weight to deaths at younger ages, and the pattern of variation is mostly dictated by the shape of the age weighting function as PALYs is decreased when the disease starts in the very early years of life or in the older ages of life with a short duration[15]. Therefore, the most accurate PALYs results are that which include both social values of discounting and age-weighting.

From our analysis, considering age weighting and discounting, approximately 35% of productivity years of a cow of its life span are lost due to ECF. In the case of bulls, approximately 49% of a bull’s productivity years are lost due to ECF. For oxen, approximately 35% of productivity years of oxen are lost due to ECF disease. However, introducing effective ECF control measures such as Immunization and tick control on cows, bulls, and oxen will reduce the loss of productivity years to approximately 0.02% (less than 1%), 0.01% (less than 1%), and 0.001% (less than 1%) respectively. Therefore, providing necessary resources to encourage farmers to take tick control measures (dipping and spraying cattle with acaricides) and routine Immunization will improve cattle productivity and lessen the disease burden of ECF and other tick-borne diseases. However, the challenge faced by these farmers is the inadequate supply of acaricides within Namwala, which results in farmers incurring extra cost to travel to Choma, the nearest district, to purchase acaricides. Also, farmers in the areas of Maala and Itezhi tezhi have no access to the Chitongo vaccine for the Immunization of their cattle, thereby resulting in a high number of ECF cases, particularly in Itezhi-tezhi therefore, farmers come to Namwala central veterinary district office to have their cattle sprayed and treated.

### Conclusion and Recommendations

The value of 517,165.40 PALYs represents the loss of healthy years of life and loss of quality of life for cattle due to ECF. The larger the number, the more the loss in cattle productivity, in this case, cows, bulls, and oxen, which are used in the farmers’ major socio-economic activities. Consequently, high PALYs are an indication of potential economic losses to livestock farmers. East Coast fever is classified as a management disease and not a disease of national economic importance. Assessing PALYs for all animal diseases will help reclassify animal diseases based on their societal burden and not only economic assessment. PALYs calculations are helpful in cost-effective analysis, in particular, comparing different health intervention programs for the same disease. PALYs are a tool in health policy that translates epidemiological data into useful information for decision-making. Based on the study findings, there is a need for further research on estimating the societal burden of all animal diseases countrywide using PALYs to aid in assessing which disease needs prioritization to minimize the loss of cattle productivity through morbidity and mortality. The results would not make more sense for decision-making on disease classification of national economic importance without ranking all animal diseases using PALYs. This study forms a basis for ranking all diseases and has an accurate measure of animal diseases’ societal burden.

## Acknowledgments

We would like to thank Mr. Ladislav Moonga for assisting with data collection and all the respondents to the questionnaires for making this study possible. The research work was supported by a grant from the National Institute of Allergy and Infectious Diseases of the National Institutes of Health ‘Spatial eco-epidemiology of tick-borne rickettsial pathogens’ under award number R01AI136035 [PI – Gaff H], through a sub-award to the University of Zambia [PI – Chitanga S]. The content is solely the responsibility of the authors and does not necessarily represent the views of the National Institutes of Health.

